# Alpha peak frequency is linked to episodic memory impairment in pathological aging

**DOI:** 10.1101/2021.06.18.448931

**Authors:** Delphine Puttaert, Vincent Wens, Patrick Fery, Antonin Rovai, Nicola Trotta, Nicolas Coquelet, Sandra De Breucker, Niloufar Sadeghi, Tim Coolen, Serge Goldman, Philippe Peigneux, Jean-Christophe Bier, Xavier De Tiège

**Affiliations:** Laboratoire de Cartographie fonctionnelle du Cerveau (LCFC), UNI – ULB Neuroscience Institute, Université libre de Bruxelles (ULB), Brussels, Belgium; Neuropsychology and Functional Neuroimaging Research Unit (UR2NF), Center for Research in Cognition and Neurosciences (CRCN), UNI – ULB Neuroscience Institute, Université libre de Bruxelles (ULB), Brussels, Belgium; Clinic of Functional Neuroimaging, Service of Nuclear Medicine, CUB Hôpital Erasme, Université libre de Bruxelles (ULB), Brussels, Belgium; Service of Neuropsychology and Speech Therapy, CUB Hôpital Erasme, Université libre de Bruxelles (ULB), Brussels, Belgium; Department of Geriatrics, CUB Hôpital Erasme, Université libre de Bruxelles (ULB), Brussels, Belgium; Department of Neurology, CUB Hôpital Erasme, Université libre de Bruxelles (ULB), Brussels, Belgium; Department of Radiology, CUB Hôpital Erasme, Université libre de Bruxelles (ULB), Brussels, Belgium

**Author notes:** ***Corresponding author:*** Delphine Puttaert.

**Keywords:** episodic verbal memory, declarative memory, FCSRT, MEG, alpha peak frequency, alpha relative power

## Abstract

The Free and Cued Selective Reminding Test (FCSRT) is a largely validated neuropsychological test for the identification of amnestic syndrome from the early stage of Alzheimer’s disease (AD). Previous electrophysiological data suggested a slowing down of the alpha rhythm in the AD-continuum as well as a key role of this rhythmic brain activity for episodic memory processes. This study therefore investigates the link between alpha brain activity and alterations in episodic memory as assessed by the FCSRT. For that purpose, 37 patients with altered FCSRT performance underwent a comprehensive neuropsychological assessment, supplemented by ^18^F-fluorodeoxyglucose positron emission tomography/structural magnetic resonance imaging (^18^FDG-PET/MR), and 10 minutes of resting-state magnetoencephalography (MEG). The individual alpha peak frequency (APF) in MEG resting-state data was positively correlated with patients’ encoding efficiency as well as with the efficacy of semantic cues in facilitating patients’ retrieval of previous stored word. The APF also correlated positively with patients’ hippocampal volume and their regional glucose consumption in the posterior cingulate cortex. Overall, this study demonstrates that alterations in the ability to learn and store new information for a relatively short-term period are related to a slowing down of alpha rhythmic activity, possibly due to altered interactions in the extended mnemonic system. As such, a decreased APF may be considered as an electrophysiological correlate of short-term episodic memory dysfunction accompanying pathological aging.

## 1 Introduction

Worldwide, about 50 million individuals are affected by dementia whose main cause is Alzheimer’s disease (AD), accounting for ∼60-70% of all dementia cases (World Alzheimer Report, 2018).

People at risk of developing dementia include patients with mild cognitive impairment (MCI) who are cognitively impaired but still independent for everyday activities (Albert et al., 2011). MCI may stem from different etiologies (e.g., degenerative, vascular, etc.). People with amnestic MCI (aMCI), which is mainly characterized by episodic memory impairment, are more likely to develop typical Alzheimer’s clinical syndrome (ACS) compared with those with non-amnestic MCI, which is characterized by other kinds of cognitive impairments (Petersen and Negash, 2008).

Clinically, AD-related predementia and dementia stages are typically characterized by an early amnestic syndrome of the hippocampal type (Dubois & Albert, 2004), which is usually evidenced using the Free and Cued Selective Reminding Test (FCSRT; Grober et al., 1988; Van der Linden et al., 2004). The FCSRT allows to investigate the individual performance in the successive steps of memory formation (i.e., encoding, storage and retrieval) in a single episodic memory task (Van der Linden et al., 2004). The first cognitive symptoms to occur in aMCI and typical AD are specific impairments in the encoding or storage of new information. These deficits are optimally revealed with memory tests that enhance mnemonic retrieval processes by means of promoting encoding specificity, such as the FCSRT (Peña-Casanova, 2012). Recent studies have demonstrated that FCSRT has a satisfying discriminative power between different forms of dementia (Teichmann et al., 2017), and is a predictor of the conversion from MCI to typical form of ACS (Grande et al., 2018).

The neural correlates of FCSRT performance in the AD-continuum have been previously identified using different neuroimaging techniques. Structural magnetic resonance imaging (sMRI) showed that higher encoding and storage scores are associated with thicker cortices in the entorhinal and parahippocampal regions, whereas a higher retrieval index is related to thicker cortices in a widespread network including the frontal regions, probably due to the role of executive functions in the active recollection of previously stored information (Epelbaum et al., 2018). Furthermore, higher scores for the three processes have been associated with a larger hippocampal volume (Epelbaum et al., 2018). Functional MRI (fMRI) demonstrated activity changes in the left inferior parietal and left precuneus to positively correlate with retrieval performance scores (McLaren et al., 2012). Neurometabolic correlates of FCSRT, as assessed through positron emission tomography using [^18^F]-fluorodeoxyglucose (FDG-PET), were also identified (Epelbaum et al., 2018). Encoding and storage scores correlated positively with glucose metabolism levels in posterior cingulate gyri, temporoparietal junctions and medial temporal lobes. Conversely, the retrieval index was positively correlated with metabolic measures in a more diffuse network including prefrontal, parietal and temporal brain regions. These cerebral regions associated with FCSRT performance closely match the neuroanatomic distribution of the neuropathological hallmarks of AD (i.e., extracellular Aβ plaques and intracellular neurofibrillary tangles) that mainly encompass the default-mode network (DMN; Canter et al., 2016). The DMN is a set of brain regions (i.e., precuneus, posterior cingulate cortex, medial prefrontal cortex, temporoparietal junctions and hippocampus) in which activity is higher when the mind is engaged in spontaneous cognition and lower during focused attention on the external environment or during goal-directed tasks (Buckner et al., 2008).

Electrophysiological techniques such as electroencephalography (EEG) and magnetoencephalography (MEG) have also been found particularly relevant in the evaluation of pathological aging and its associated cognitive symptoms (for a review, see Maestú et al., 2019). They have consistently highlighted a spectral correlate of AD, i.e., a decrease in both the alpha peak frequency (APF) and relative power compared to healthy elders (for a review, see López-Sanz et al., 2019). A reduction of the APF has indeed been repeatedly observed in AD (e.g., from 9.4 Hz to 8.0 Hz in Poza et al., 2007; see also Passero et al., 1995; Montez et al., 2009; Babiloni et al., 2017) and, to a lesser extent, in MCI (e.g., from 9.8 Hz to 9.0 Hz in López-Sanz et al., 2016; see also Garcés et al., 2013). A decrease in alpha power was also reported in both of these symptomatic groups (Passero et al., 1995; Huang et al., 2000; Moretti et al., 2004; Engels et al., 2016; López-Sanz et al., 2016; Babiloni et al., 2017; Koelewijn et al., 2017). The decrease in APF further correlated with a reduction in hippocampal volume (Garcés et al., 2013; López-Sanz et al., 2016), a well-demonstrated structural damage in AD (Henneman et al., 2009). All these data suggest that a slowing of the alpha rhythmic activity could take place early in the symptomatic AD-continuum, potentially due to alterations in thalamo-cortical interactions secondary to, e.g., loss of thalamic inhibitory interneurons (Bhattacharya et al., 2011). Given how episodic memory is affected along the AD-continuum, these results are also in line with electrophysiological studies that demonstrated the key role of rhythmic brain activity in the alpha frequency band for episodic memory processes (Klimesch et al., 1993; Klimesch, 1999; Hanslmayr et al., 2016; Sutterer et al., 2019; Martín-Buro et al., 2020).

To the best of our knowledge, no study has yet investigated the link between spectral parameters (i.e., APF and relative power) of the alpha rhythmic brain activity (henceforth referred to as *“alpha activity”*) and alterations in episodic memory processes as assessed by the FCSRT. The present study therefore aims at closing this gap by using MEG recordings obtained in a group of elders (henceforth referred to as “*patients*”) exhibiting altered FCSRT performance in the encoding or storage processes. More specifically, we first compared their neuropsychological performance, hippocampal volume, regional cerebral glucose metabolism and spectral parameters (i.e., APF and relative power) of their alpha activity with those of matched healthy elders with preserved FCSRT performance to characterize their overall clinical profile. We then searched in patients for associations linking these spectral parameters to FCSRT scores, hippocampal volume, and regional cerebral glucose metabolism. Given the reported alterations in alpha activity along the AD-continuum and the key role of alpha activity in episodic memory processes, we expected (i) to find strong associations between its spectral parameters and FCSRT performance, and (ii) that variations in its spectral parameters will be related with hippocampal volume and regional cerebral glucose metabolism.

## 2 Materials and Methods

### 2.1 Participants

From February 15, 2018 to February 27, 2020, we included in this study 37 elderly patients who benefited from a comprehensive neuropsychological evaluation at the consultation of D.P. at the CUB Hôpital Erasme, based on the following inclusion criteria: (i) age range 55– 90 years, (ii) significant alterations in encoding or storage of new information as evidenced with the FCSRT (for more details, see *Clinical and neuropsychological evaluation* and *Diagnostic criteria* below), (iii) no contraindication for MEG and PET-MR investigations (e.g., claustrophobia, ferrous metal implants, etc.), (iv) written informed consent to contribute to the study, and (v) absence of exclusion criterion. Exclusion criteria included: a prior history of neurological or psychiatric disorder, a chronic use of psychotropic drugs or alcohol, insufficient level in French language, severe visual or hearing impairment, and a modified Hachinski score ≥ 4 (Hachinski Ischemic Score; Hachinski, 1992). Nineteen healthy elders were also recruited for comparison purposes based on similar inclusion criteria except that they performed within the normal range at the FCSRT.

All individuals had a least 6 years of education and almost all of them were right-handed (3 ambidextrous and 3 left-handed) according to the Edinburgh inventory (Oldfield, 1971).

The study protocol was approved by the Ethics Committee of the CUB Hôpital Erasme (P2017/427, Brussels, Belgium). All experiments were performed in accordance with the standards of the Good Clinical Practice and the Declaration of Helsinki.

### 2.2 Clinical and neuropsychological evaluation

The comprehensive clinical and neuropsychological evaluation has been described in detail elsewhere (Puttaert et al., 2020).

Briefly, the clinical evaluation included the mini mental state examination (MMSE; Folstein et al., 1983), the neuropsychiatric inventory (NPI; Cummings, 1997), the short form of the geriatric depression scale (GDS-15; Yesavage et al., 1982), the clinical dementia rating (CDR; O’Bryant et al., 2008), and the basic and instrumental activities of daily living (bADLs and iADLs) questionnaires (Lawton and Brody, 1969). The neuropsychological assessment comprised the evaluation of verbal (Van der Linden et al., 2004) and visual (Baddeley et al., 1994) episodic memory, working memory (Wechsler Memory Scale, WMS-III), language abilities (Bachy Langedock, 1988), executive functions (Reitan, 1958; Henry et al., 2004; Hutchison et al., 2010), and visuo-constructive abilities (Berry et al., 1991).

#### FCSRT-derived learning processes indices

Each participant underwent the French version of the FCSRT to specifically evaluate the integrity of the encoding, storage and retrieval steps in verbal episodic memory formation (Grober et al., 1988; Van der Linden et al., 2004).

Briefly, in the FCSRT, participants were instructed to learn a list of 16 words presented 4 by 4 on sheets and semantically related with a cue (e.g., “flower” is the cue for the item “daffodil”). First, they had to point and name the words corresponding to the semantic cue provided by the experimenter, and then immediately recall words when presented with the cue (Immediate Recall; IR). Second, they had to freely recall as many words as possible in 2 minutes (Free Recall; FR) and a semantic cue was provided for the missing ones (Cued Recall; CR). The Total Recall (TR) score added FR and CR scores (i.e., the number of words correctly recalled). In total, three recall, first free (FR1, FR2 and FR3) and then cued, trials were successively performed and were separated from each other by a distractive task (counting backwards for 20 s). After 20 minutes and after 1 week, a Delayed Recall (DR) test was administered following the same procedure. Based on this procedure, we derived the following FCSRT indices:

- **Encoding impairment index**, defined as the number of words that were not recalled in FR1, those neither recalled in FR1 nor in FR2, and those never recalled.
- **Index of Sensitivity to Cueing** (ISC, Epelbaum et al., 2011), computed as 100 * (sum of the three cued recall / (48 – sum of 3 FR)). This ISC index provides an estimate of the magnitude of sensitivity to semantic cues and was previously proposed to be a specific marker of hippocampal-related memory (Dubois et al., 2007).
- **Long-term retention rate index**, defined as the ratio of the total DR score after 1 week to the total DR score after 20 min. This index measures long-term (1 week) memory storage performance.

FCSRT scores below 2 standard deviation (SD) of the appropriate normative mean (Van der Linden et al., 2004) were considered as abnormal.

### 2.3 Diagnostic criteria

Healthy elders performed normally at the comprehensive neuropsychological assessment and had no subjective memory complaint (evaluated here with the GDS-short form question: “*Do you feel you have more problems with your memory than most?*”; Yesavage et al., 1982).

Patients with altered performance at the FCSRT were classified as aMCI if they had (i) a subjective and objective cognitive impairment as described by the National Institute on Aging – Alzheimer’s Association (NIA-AA) workgroups criteria (Albert et al., 2011), specifically affecting episodic memory (Petersen et al., 1999), and (ii) a normal functioning in their daily life activities (IADL, Lawton & Brody, 1969).

Patients with altered FCSRT performance were classified as having ACS (i.e., clinically ascertained multi or single domain amnestic syndrome) if they fulfilled the new criteria of NIA-AA (Jack et al., 2018) including the evident functional impact on daily life.

### 2.4 Neuroimaging investigations

All participants (i.e., healthy elders and patients) underwent a comprehensive structural and functional neuroimaging investigation that included resting-state MEG prior to hybrid brain PET-MR scanning comprising simultaneous optimized sMRI and FDG-PET. Both MEG and PET-MR investigations were performed on the same day in all participants.

#### MEG data acquisition, preprocessing and spectral parameters estimation

Neuromagnetic data were acquired in a quiet resting-state during two 5-min sessions (sitting position, eyes open with fixation cross, online band-pass filter : 0.1–330 Hz, sampling rate : 1000 Hz) using a 306 channel whole-scalp MEG system (Triux, MEGIN, Croton Healthcare, Helsinki, Finland) placed inside a light-weight magnetically shielded room (Maxshield, MEGIN, Croton Healthcare, Helsinki, Finland, see De Tiège et al., 2008 for more details) at the CUB Hôpital Erasme (Brussels, Belgium). The position of participants’ head was recorded continuously inside the MEG helmet with four head tracking coils. Their location was localized relative to anatomical fiducials with an electromagnetic tracker (Fastrak, Polhemus, Colchester, Vermont, USA).

The first step of MEG data preprocessing consisted in the application of the spatiotemporal extension of signal space separation (correlation coefficient: 0.98; window length: 10 s) in order to remove external interferences and nearby sources of magnetic artifact, and also compensate for head movements (Taulu and Simola, 2006). In a second step, remnant cardiac, ocular and system artifacts were visually identified and eliminated using an independent component analysis (FastICA algorithm with dimension reduction to 30 components, hyperbolic tangent nonlinearity function, Vigario et al., 2000) of the filtered data (off-line band-pass filter: 0.1–45 Hz). Only magnetometers data were used in the subsequent analysis as done in López-Sanz et al., 2016.

The power spectrum of cleaned MEG signals was then estimated at 0.1 Hz spectral resolution as the magnitude square of Fourier coefficients obtained via finite Fourier transformation over 10 s-long non-overlapping epochs and averaged over these epochs. This spectral data was further averaged over temporo-occipital sensors to yield a single spectral profile per participant. The APF and its relative amplitude were then identified individually by fitting a model spectrum composed of (i) a Gaussian peak that aims at detecting the alpha peak superposed to (ii) a power law reflecting the background noise and other signals of no interest. More specifically, we used a two-step procedure inspired by López-Sanz et al. (2016). First, a power law was fitted to the MEG spectrum within a frequency domain well outside the classic alpha frequency band and comprising 1–4 Hz and 20–30 Hz. This power law was then evaluated on the whole frequency domain and subtracted from the MEG spectrum to eliminate typical low-frequency power increases. Second, a Gaussian bell curve (not normalized to unit integral) was fitted to this corrected MEG spectrum within the 7–13 Hz frequency range covering the alpha band. The estimated Gaussian mean was taken as a measure of the APF. The relative amplitude at this frequency was taken as the ratio of the APF power of full spectral model (sum of the power law and Gaussian curves), to that of the background power law only.

#### Hybrid ^18^F-FDG-PET/MR data acquisition and preprocessing

A T1-weighted brain MRI (Repetition Time/Echo Time/Flip Angle: 8.3 ms/3.1 ms/12°, TI: 450 ms, field of view (FOV): 24 cm x 24 cm, matrix: 240 × 240, resolution: 1 mm × 1 mm × 1 mm) and brain FDG-PET were acquired simultaneously on a hybrid 3 T SIGNA PET-MR scanner (GE Healthcare, Milwaukee, Wisconsin, USA) at the CUB Hôpital Erasme (Brussels, Belgium).

The FreeSurfer software (version 6.0; Martinos Center for Biomedical Imaging, Massachusetts, USA) and the Desikan-Killiany atlas (Desikan et al., 2006) were used to obtain the estimated-total intracranial volume (eTIV) and the whole hippocampal volume from sMRI. The whole hippocampal volume was calculated as the sum of left and right whole hippocampal grey matter volumes.

The preprocessing of FDG-PET images was performed with SPM8 (www.fil.ion.ucl.ac.uk/spm) and spatially normalized using a specific FDG-PET aging and dementia template (Della Rosa et al., 2014), which was itself normalized to the Montreal Neurological Institute (MNI) template. Data were then smoothed with a FWHM 8-mm Gaussian kernel. The photons’ attenuation specific of the PET acquisition was corrected with a MRI-based map (MRAC) acquired simultaneously. PET images were displayed in a 256 × 256 × 89 matrix format, with a slice thickness of 2.78 mm. The reconstructed files were downloaded in their original format (DICOM, ECAT, Interfile) for meta-information and converted in NIfTI format for further analysis.

### 2.5 Statistical analyses

#### Differences between healthy elders and patients with altered FCSRT

Firstly, we estimated the group differences in terms of behavior (i.e., FCSRT scores), regional structural (i.e., whole hippocampal volume) and metabolic (i.e., regional glucose metabolism) brain data, and spectral parameters of the alpha activity (APF and relative power) between healthy elders and patients. The main goal of these analyses was to characterize the population of patients with altered FCSRT performance relative to healthy elders.

Between-group differences in episodic memory indices were evaluated with an ANCOVA analysis (*p* < .05 Bonferroni corrected for the number of indices considered) controlling for age, years of education and gender (used as covariates of no interest).

Differences in whole hippocampal volume (relative to eTIV) were computed using an ANCOVA analysis (*p* < .05 Bonferroni corrected for age, years of education and gender). The whole hippocampal volume was represented here as a percentage of eTIV in order to adjust for differences in head size across participants. Our focus on the hippocampus was driven by prior reports indicating that hippocampal volume is a structural correlate of FCSRT performance (e.g., Albert et al., 2011).

Subtractive voxel-based analysis of FDG-PET data were also performed as previously done in previous studies from our group (De Tiege et al., 2004; De Tiège et al., 2008b; Puttaert et al., 2020). We used SPM12 (http://www.fil.ion.ucl.ac.uk/spm/,Wellcome Trust Centre for Neuroimaging, London, UK) to construct general linear models (GLMs) of the preprocessed FDG-PET data of healthy elders and patients with altered FCSRT performance taken as separate groups. Age, gender and years of education were introduced as covariates of no-interest. Proportional scaling was applied beforehand to remove inter-subject variation in global brain metabolism and an explicit FDG mask was used to restrict the analyses inside the brain. Separate mass-univariate *T* contrasts then searched, throughout the brain, for regions showing significant decrease or increase in metabolism between healthy elders and patients. Results were considered significant at *p* < .05 family-wise error (FWE)-corrected unless otherwise stated.

Finally, differences across groups regarding spectral parameters of the alpha activity were assessed using an ANCOVA procedure and Bonferroni *post hoc* analyses with age, gender and education as covariates of no-interest.

#### Correlation analyses in patients with altered FCSRT

Pearson’s correlation analysis was performed in patients with altered FCSRT performance only in order to explore the relationship between electrophysiological data (i.e., APF and relative power) and FCSRT indices (see 2.2), structural (i.e., whole hippocampal volume) and metabolic (i.e., regional glucose metabolism) brain data. The covariates of no-interest used in these analyses were similar to those presented before for the group-difference analyses (i.e., age, gender and years of education).

The results of the correlation between electrophysiological data and each FCSRT index were considered significant at *p* < .05 Bonferroni corrected for multiple comparisons following the number of FCSRT indices considered. A similar Pearson correlation analysis was then performed with the whole hippocampal volume.

Finally, a mass-univariate version of Pearson correlation analysis was used to investigate the relationship between spectral parameters of the alpha activity and regional cerebral glucose metabolism. We used SPM12 (http://www.fil.ion.ucl.ac.uk/spm/, Wellcome Trust Centre for Neuroimaging, London, UK) to construct two (i.e., one for APF and one for alpha relative power) separate GLMs of the preprocessed patients’ FDG-PET data (i.e., 37 scans), each including one of the two spectral parameters of the alpha activity as a covariate of interest. Age, gender and years of education were used as covariates of no-interest. Proportional scaling was applied to remove inter-subject variations in global brain metabolism and an explicit FDG mask was used to restrict the analyses inside the brain. Separate *T* contrast analyses then searched, throughout the brain, for regions showing significant positive or negative correlations with spectral parameters of the alpha activity. Results were considered significant at *p* < 0.05 corrected for multiple comparisons (FWE rate) at the voxel level or at p_FWE_ < 0.05 at the cluster level (height threshold: *p* < 0.001).

## 3 Results

### 3.1 Participants

#### Demographic and clinical data

The included participants comprised 19 healthy elders (mean age and standard deviation: 64.31 ± 7.60 years, age-range: 55-78 years, 15 females) with FCSRT performance in the normal range, and 37 patients with altered FCSRT performance (73.08 ± 7.30 years, age-range: 56-92 years, 33 females).

Among the 37 patients, 19 were clinically classified as aMCI (71.22 ± 5.9 years; 12 females) and 18 patients as ACS (74.94 ± 8.22 years; 11 females). The aMCI and ACS patients were comparable in age, gender, years of education, depression status and laterality.

The characteristics of all participants are shown in Table 1.

**Table 1.** Participants’ demographic and clinical data for healthy elders (HE) and patients. Values are presented as mean ± SD (standard deviation). HE: healthy elders, aMCI: amnestic-mild cognitive impairment, ACS: Alzheimer’s clinical syndrome, GDS: Geriatric Depression Scale (short form), CDR: Clinical Dementia Rating scale, sMRI: structural magnetic resonance imaging, WHV: whole hippocampal volume, eTIV: estimated-total intracranial volume, APF: alpha peak frequency, ARP: alpha relative power, MMSE: Mini Mental State Examination, FCSRT: Free and Cued Selective Reminding Test, LT: long-term, ISC: index of sensitivity to cueing, FR: free recall, TR: total recall, DR: delayed recall. Group differences were assessed with *χ*^2^ (Pearson) for categorized indices, independent samples *t*-test for continuous variables normally distributed and with a Mann-Whitney test for variables not normally distributed.

|  | Controls | Patients |  | *p* value |  |
| --- | --- | --- | --- | --- | --- |
|  | HE<br>(n = 19) | aMCI<br>(n = 19) | ACS<br>(n = 18) | HE vs.<br>patients | aMCI vs.<br>ACS |
| Age, years | 64.31 ± 7.60 | 71.22 ± 5.90 | 74.94 ± 8.22 | <.001 | .13 |
| Gender, females | 15 | 12 | 11 | .25 | .74 |
| Education, years | 14.26 ± 4.28 | 12 ± 3.80 | 12.77 ± 3.52 | .34 | .55 |
| GDS | 1.89 ± 2.20 | 3 ± 3.04 | 4.05 ± 3.73 | .03 | .37 |
| CDR | 0 ± 0 | 0.5 ± 0 | 0.7 ± 0.25 | - | - |
| **sMRI** |  |  |  |  |  |
| WHV, % eTIV | 44.3 ± 5.9 | 40.2 ± 4.9 | 33.6 ± 4.2 | .004 | <.001 |
| **Spectral parameters of alpha activity** |  |  |  |  |  |
| APF, Hz | 9.44 ± 1.26 | 9.6 ± 1.21 | 8.36 ± 1.01 | .46 | .009 |
| ARP | 3.82 ± 2.48 | 3.56 ± 1.49 | 4.88 ± 2.18 | .29 | .11 |
| **Global cognition score** |  |  |  |  |  |
| MMSE | 28.68 ± 1.20 | 25.70 ± 2.14 | 22.16 ± 5.33 | <.001 | .01 |
| **FCSRT indices and scores** |  |  |  |  |  |
| Encoding impairment | 0.05 ± 0.22 | 12.07 ± 11.02 | 27.57 ± 10.62 | <.001 | <.001 |
| LT retention rate | 88.4 ± 8.9 | 29.1 ± 27.1 | 19.7 ± 33.4 | <.001 | .38 |
| ISC | 98.41 ± 3.87 | 53.24 ± 20.74 | 27.15 ± 20.48 | <.001 | <.001 |
| Sum of FR | 35.05 ± 4.12 | 14.93 ± 7.07 | 5.52 ± 5.69 | <.001 | <.001 |
| Sum of TR | 47.73 ± 0.65 | 32.43 ± 10.65 | 16.17 ± 10.6 | <.001 | <.001 |
| Free DR after 20' | 14.10 ± 1.56 | 3.75 ± 2.49 | 1.17 ± 1.74 | <.001 | .002 |
| Total DR after 20' | 16 ± 0 | 9.75 ± 4.05 | 4.94 ± 3.61 | <.001 | - |
| Free DR after 1 week | 7.25 ± 2.81 | 1.27 ± 1.9 | 0 ± 0 | <.001 | - |
| Total DR after 1 week | 14.12 ± 1.4 | 5.27 ± 3.6 | 0.81 ± 0.75 | <.001 | <.001 |

#### Differences in FCSRT indices between healthy elders and patients

A significant difference was found between healthy elders and patients for the encoding impairment index (*p* <.001, Bonferroni corrected) with a larger number of non-encoded words for patients (mean and standard deviation: 19.82 ± 13.23) in comparison with healthy elders (0.05 ± 0.22). Furthermore, healthy elders had a higher long-term retention rate than patients (*p* <.001, Bonferroni corrected). Indeed, the proportion of words recalled after one week that were also recalled after 20 minutes was higher in healthy elders (average percentage ± SD: 88.4 % ± 8.9 %) than patients (24.3 % ± 30.4 %). Finally, the results demonstrated a significant difference (*p* <.001, Bonferroni corrected) for the ISC index with a higher score for healthy elders (average percentage ± SD: 98.41 % ± 3.87 %) than patients (40.19 % ± 24.24 %).

#### Differences in whole hippocampal volume, regional brain glucose metabolism, and spectral parameters of the alpha activity between healthy elders and patients

The whole hippocampal volume (relative to eTIV) was significantly different between healthy elders and patients (*p* = .004; Bonferroni corrected). Compared with healthy elders (mean and standard deviation: 44.3 % ± 5.9 %), patients (36.9 % ± 5.6 %) showed significant decrease in the whole hippocampal volume.

Voxel-based subtractive analyses of FDG-PET data demonstrated that patients had a significant reduction in glucose consumption in left precuneus, left inferior parietal gyrus, right angular gyrus and right inferior temporal gyrus compared with healthy elders (*p* < .05 with FWE correction and cluster threshold at *k* > 100 voxels; Figure 1).

**Figure 1.**
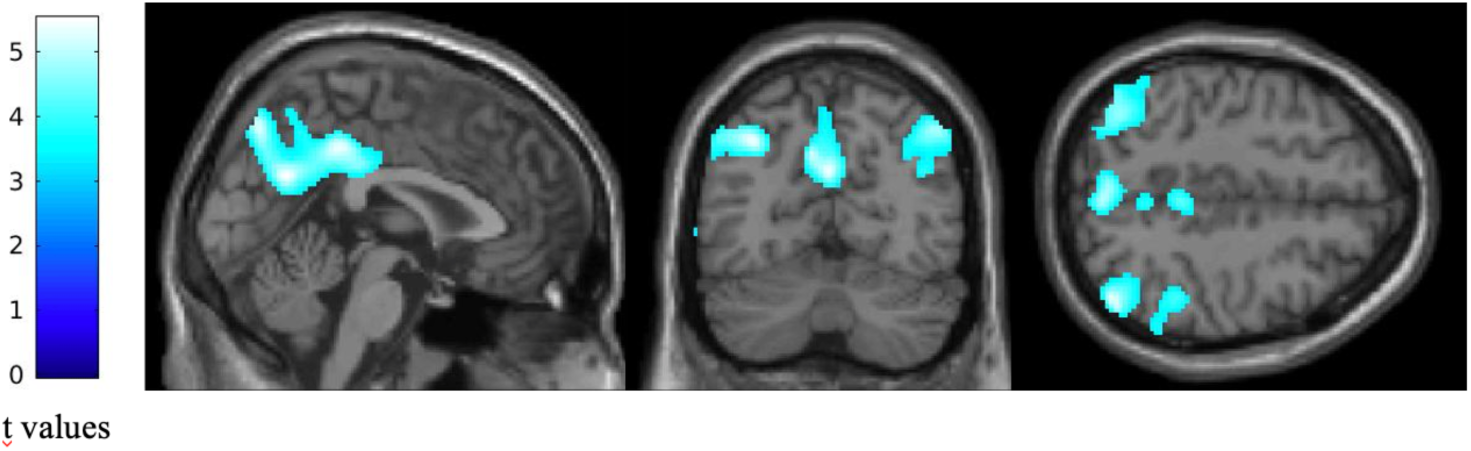
Statistical parametric *T* maps showing a significant reduction in regional glucose consumption in patients with altered FCSRT performance compared with healthy elders. Images are thresholded at *p* < .05, FWE-corrected with a cluster threshold at 100 voxels for visualization purpose. The color scales on the left represent the T-statistic of the significant voxels.

No significant difference was disclosed between healthy elders and patients for the APF (*p* = .46; Bonferroni corrected, *p* = .34; uncorrected; healthy elders, mean and standard deviation: 9.44 Hz ± 1.26 Hz; patients, 9.05 Hz ± 1.27 Hz), nor for the alpha relative power (*p* = .29; Bonferroni corrected, *p* = .62; uncorrected; healthy elders, 3.82 ± 2.48; patients, 4.15 ± 1.92).

### 3.2 Correlation analyses in patients

#### Between spectral parameters of alpha activity and FCSRT indices

Correlation analyses showed a significant negative association (r = -.54; *p*_corrected_ = .008, *p*_uncorrected_ = .009; see Figure 2) between the encoding impairment index and the APF. Indeed, the lower was the frequency of the APF, the higher was the number of non-encoded words in patients. Regarding the ISC, a significant positive correlation was observed with the APF (r = .46; *p*_corrected_ = .014, *p*_uncorrected_ = .01; see Figure 2) suggesting that a higher APF was related to a better sensitivity to cueing in order to retrieve previously stored words. No significant correlation was found with the long-term retention rate nor with the alpha relative power (all *ps* > .05, corrected).

**Figure 2.**
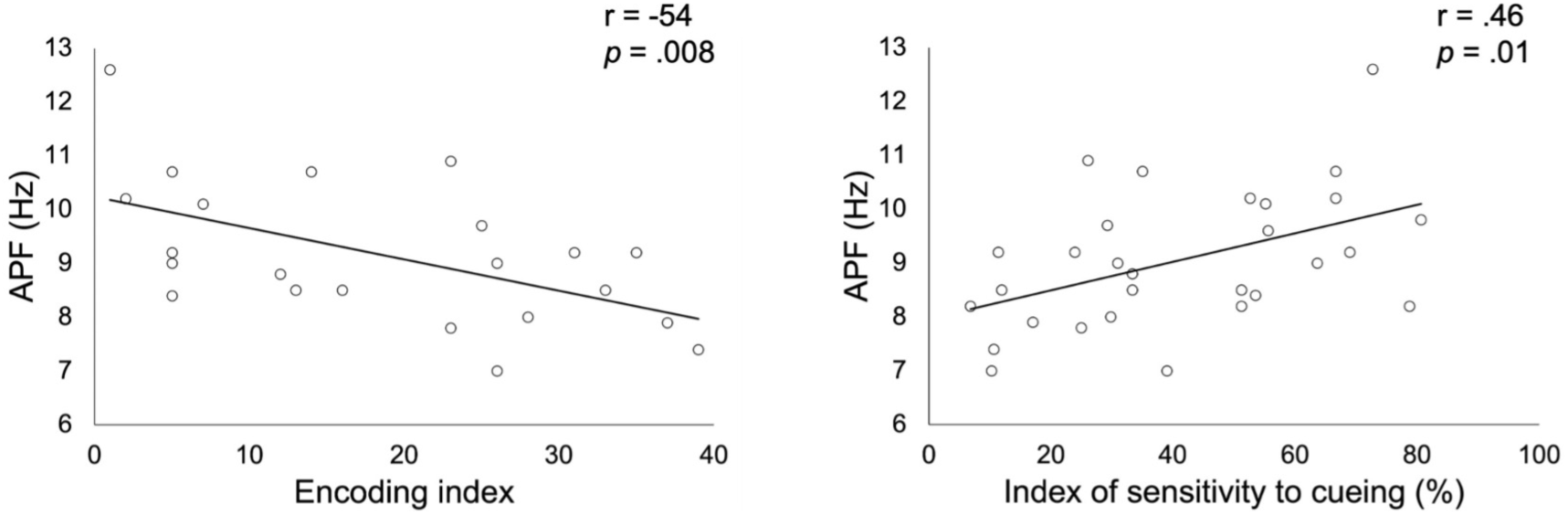
Correlation between alpha peak frequency (APF) and the encoding impairment index (left side) or the index of sensitivity to cueing (right side) obtained in patients with altered FCSRT performance.

#### Between spectral parameters of alpha activity and whole hippocampal volume

Results showed a significant positive correlation (r = .59, *p* < .001, Figure 3) between whole hippocampal volume and APF in patients. A significant negative correlation (r = -.48, *p* = .008, Figure 5) was also found between whole hippocampal volume and alpha relative power in patients altered in FCSRT.

**Figure 3.**
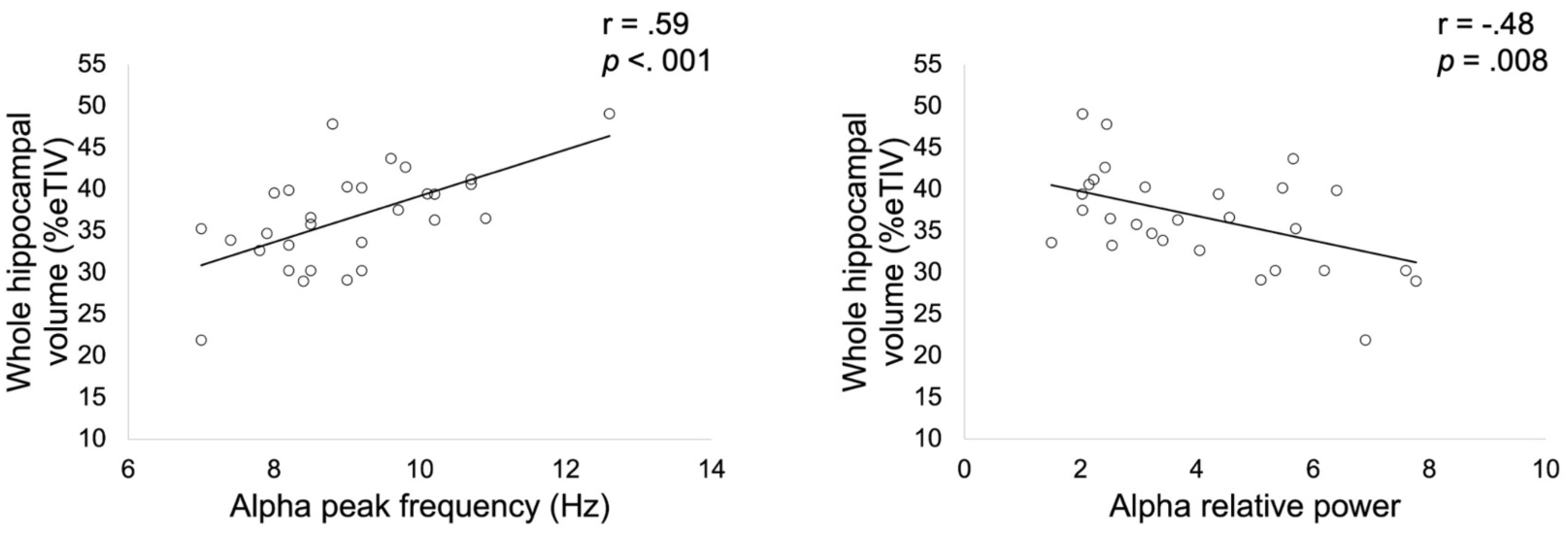
Correlation between whole hippocampal volume expressed as a percentage of total intracranial volume (TIV) and alpha peak frequency (left side) or relative alpha power (right side) obtained in patients with altered FCSRT performance.

#### Between spectral parameters of alpha activity and regional cerebral glucose metabolism

A positive correlation was observed between APF and the metabolic activity of bilateral posterior cingulate cortex (PCC; *p* _FWE_ = .05, cluster level; Figure 4).

**Figure 4.**
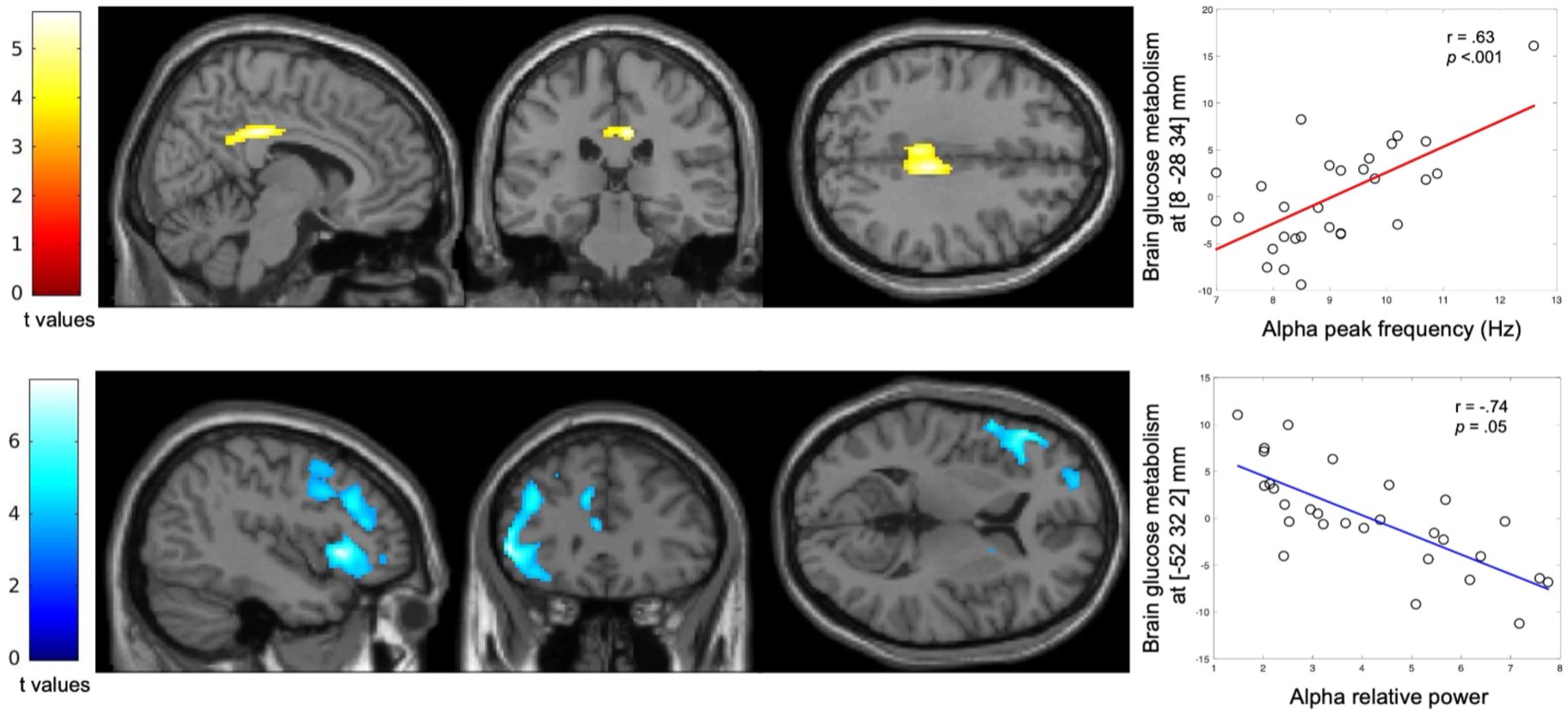
Statistical parametric T maps showing a significant positive correlation between APF and regional glucose consumption (Top) and a significant negative correlation between alpha relative power and regional glucose consumption across the patients with altered FCSRT performance. Images are thresholded at p < 0.001 uncorrected with a cluster threshold at 100 voxels for visualization purpose. The color scales represent the T-statistic of the significant voxels.

A significant negative correlation was found between alpha relative power and the glucose metabolism in the left inferior frontal gyrus (*p* _FWE_ <.001, cluster level; Figure 4).

## 4 Discussion

This multimodal neuroimaging study demonstrates in patients with altered FCSRT performance that (i) the APF is positively related to the encoding efficiency as well as to the sensitivity of cueing in verbal episodic memory, but not to the long-term retention performance, (ii) the APF is positively related to the whole hippocampal volume and the level of relative glucose metabolism of the PCC, and (iii) the alpha relative power is negatively associated with the whole hippocampal volume and the metabolic activity in the left frontal gyrus.

This study reveals that the APF in patients with altered FCSRT performance is related to alterations in the ability to learn and store new information for a relatively short period of time (i.e., within minutes or 1-2 hours), but not in the long-term (i.e., 1 week). This finding is compatible with a previous MEG study that found a positive relation between the APF and short-term storage (i.e., immediate and delayed recall scores in Weschler memory scale (WMS-III)) in verbal memory (López-Sanz et al., 2016). It is also perfectly in line with the significant association between the APF and the whole hippocampal volume found in this study, but also in previous MEG studies (Garcés et al., 2013; López-Sanz et al., 2016), considering the well-recognized role of the hippocampus in the first steps of episodic memory formation (i.e., encoding and storage processes) (Sarazin et al., 2010). We also found a positive correlation between the APF and the glucose metabolism in the PCC, which is increasingly recognized as playing a key role in episodic memory processes, and especially in encoding and retrieval processes (Wagner et al., 2005; Vilberg and Rugg, 2009; Lega et al., 2017; Natu et al., 2019). For example, deep brain stimulation (DBS) of the PCC during episodic memory encoding has been shown to impair subsequent recall, highlighting its implication in episodic memory encoding (Natu et al., 2019).

A decrease in the APF has been previously demonstrated in pathological aging such as in MCI (Garcés et al., 2013; López-Sanz et al., 2016) and AD (Passero et al., 1995; Montez et al., 2009). Different hypotheses have been proposed to explain this slowing of the alpha activity observed in pathological aging but its underlying neural cause is still not clearly understood. Neural network simulation studies have previously demonstrated that alterations in the dynamics of cortico-thalamic circuits could account for the decrease of the APF observed in patients with MCI or AD (Bhattacharya et al., 2011; Hindriks and van Putten, 2013; Li et al., 2020). Interestingly, a reduction in the volume of thalamus has been previously reported in AD (Callen et al., 2001; Zarei et al., 2010; Low et al., 2019) and was associated with impaired cognitive functioning (de Jong et al., 2008). Furthermore, neurofibrillary tangle deposits were also found in the thalamus at the same Braak stage as those found in the hippocampal region (Braak and Braak, 1991). Still, it is unclear if these thalamic abnormalities are primary related to AD pathology or secondary to cortico-thalamic/hippocampo-thalamic denervation due to neuronal loss (Abuhassan et al., 2014; Aggleton et al., 2016; Jagirdar and Chin, 2020).

Both the PCC and the thalamus, that are key components of the hippocampal network of the limbic system or Papez circuit (Catani et al., 2013; Rolls, 2015), have direct connections with the hippocampus (for a review, see Aggleton, 2012). A previous fMRI study showed a significant association between PCC and hippocampal activations during episodic memory encoding and recognition in MCI patients (Papma et al., 2017). More specifically, the limbic system is composed of three distinct networks including the hippocampal-diencephalic limbic circuit (Catani et al., 2013) that is dedicated to memory (Aggleton, 2008). Previous imaging studies have demonstrated altered metabolism and reduced functional activation of this network in neurodegenerative disorders as MCI (Minoshima et al., 1997) and AD (Buckner, 2005).

Taken together, these data and results of the present study suggest that APF decreases in pathological aging might reflect altered hippocampo-thalamo-cortical interactions leading to a reduced ability to learn and store new information in the short term (i.e., within minutes or 1-2 hours). This hypothesis is in line with the idea that memory loss in pathological aging would be related to a more widespread mnemonic circuit pathology rather than to a predominant medial temporal lobe dysfunction (for a detailed argumentation, see Aggleton et al., 2016).

Finally, a negative relation was also found between alpha relative power and metabolic activity in the left inferior frontal gyrus in patients with altered FCSRT. This finding might represent a neural correlate of the potential protective role of the prefrontal cortex in pathological aging. Indeed, a previous resting-state fMRI study showed that higher activity in left insula and left inferior frontal gyrus reduces the negative impact of AD-associated pathology on episodic memory in elders at risk for AD (Lin et al., 2016).

Of note, we did not find any significant difference in APF values (nor in alpha relative power) between healthy elders and patients with altered FCSRT. A possible explanation could be that patients had a relatively high mean MMSE score (i.e., 23.88 ± 4.43), suggesting moderate alterations in general cognitive functioning. In line with this hypothesis, a previous MEG study failed to find any significant difference in APF between healthy elders and AD patients with an even lower mean MMSE score (i.e., 20.8 ± 4.0) (Osipova et al., 2005). It could also be related to the methods chosen to calculate the APF, which can substantially vary and potentially explain some inconsistencies between studies (e.g., Osipova et al., 2005; Garcés et al., 2013; López-Sanz et al., 2016). Therefore, a clear consensus on the optimal method used to estimate the individual APF is needed in order to avoid bias and inconsistencies in the results reported in the literature. In that context, a free and open-source software has been developed to automatically calculate the APF in resting-state EEG data (Corcoran et al., 2018). This open-source software should be easily transposable to MEG data and could be useful for future studies.

The present study suffers from several limitations. First, our sample size was relatively modest and may have impacted the sensitivity to detect other relationships between neural and behavioral data (e.g., the correlation between the alpha relative power and FCSRT scores). Second, our population was mainly defined on the basis of scores obtained in the FCSRT for which a low performance can be related to various neurodegenerative diseases associated to pathological aging such as AD, frontotemporal dementia or Lewy body disease (Bertoux et al., 2014; Lemos et al., 2015; Bussè et al., 2018). For example, despite the very high sensitivity of FCSRT (e.g., 100% in Teichmann et al., 2017) for identifying typical AD patients, abnormal scores in FCSRT do not have an absolute specificity (e.g., 75% in Teichmann et al., 2017) for typical AD. Still, our patients showed a significant decrease in Encoding and Storage of new information, in whole hippocampal volume, as well as in glucose consumption in posterior temporo-parietal regions compared with healthy elders. These findings are closely similar to the “AD-like” patterns identified in the literature (see Henneman et al., 2009; Herholz et al., 2002; Sarazin et al., 2007). Third, multiple neurological and psychiatric conditions were previously associated with a lowering of the APF such as, e.g., burnout syndrome (van Luijtelaar et al., 2010), cerebral ischemia (Van der Worp et al., 1991) or cancer-related fatigue (Zimmer et al., 2015). Therefore, a slowing of alpha activity cannot be considered as pathognomonic of pathological aging (López-Sanz et al., 2019). Finally, our study was limited by its cross-sectional design and longitudinal investigations are needed to explore how the episodic memory decline, that could be observed in pathological aging (Lemos et al., 2015), can be related to the evolution of the slowing process of the alpha activity.

In conclusion, this study demonstrates that alterations in the ability to learn and store new information for a relatively short-term period of time are related to a slowing of the alpha activity, possibly due to altered neural interactions in the extended mnemonic system. As such, a decrease of the APF may be considered as an electrophysiological correlate of short-term episodic memory dysfunction accompanying pathological aging.

## Data availability

The datasets analyzed during this study are available from the corresponding author upon reasonable request and after approval by institutional authorities (i.e., CUB Hôpital Erasme and Université libre de Bruxelles).

## Author contributions

DP designed the study, collected sample and data, performed the neuropsychological assessment, performed MEG recordings, pre-processed MEG data, analyzed and interpreted behavioral and neuroimaging data, wrote the manuscript and prepared the figures. VW contributed to MEG data and statistical analyses, interpreted data and wrote the manuscript. PF designed the study, analyzed and interpreted neuropsychological data. SDB and J-CB contributed to patients’ clinical and neuroimaging data collection. AR, NS and TC contributed to MRI data acquisition and analyses. NT contributed to FDG-PET data acquisition and analyses. NC contributed to MEG data analyses. PP designed the study, interpreted behavioral and neuroimaging data and wrote the manuscript. SG designed the study, analyzed and interpreted behavioral and neuroimaging data, wrote the manuscript. XDT supervised and designed the study, analyzed and interpreted data, wrote the manuscript and obtained funding. All authors drafted the manuscript for intellectual content and approved the final version.

## Acknowledgments

Delphine Puttaert is supported by a research grant of the Fonds Erasme (research convention “Alzheimer” Brussels, Belgium). Tim Coolen is Clinical Master Specialist Applicant to a PhD and Xavier De Tiège is Postdoctoral Clinical Master Specialist at the Fonds de la Recherche Scientifique (FRS-FNRS, Brussels, Belgium).

This study has been supported by a research grant of the Fonds Erasme (research convention “Alzheimer” Brussels, Belgium).

The PET-MR project at the CUB Hôpital Erasme and Université libre de Bruxelles is financially supported by the Association Vinçotte Nuclear (AVN, Brussels, Belgium).

The MEG project at the CUB Hôpital Erasme is financially supported by the Fonds Erasme (research convention “Les Voies du Savoir”, Brussels, Belgium).

